# Production and characterisation of stabilised PV-3 virus-like particles using *Pichia pastoris*

**DOI:** 10.1101/2022.09.16.508282

**Authors:** Lee Sherry, Keith Grehan, Jessica J. Swanson, Mohammad W. Bahar, Claudine Porta, Elizabeth E. Fry, David I. Stuart, David J. Rowlands, Nicola J. Stonehouse

**Author notes:** Address correspondence to David J. Rowlands, and Nicola J. Stonehouse.

## Abstract

Following the success of global vaccination programmes using the live-attenuated oral and inactivated poliovirus vaccines (OPV and IPV), wild poliovirus (PV) is now only endemic in Afghanistan and Pakistan. However, the continued use of these vaccines poses potential risks to the eradication of PV. The production of recombinant PV virus-like particles (VLPs), which lack the viral genome offer great potential as next-generation vaccines for the post-polio world. We have previously reported production of PV VLPs using *Pichia pastoris*, however, these VLPs were in the non-native conformation (C Ag), which would not produce effective protection against PV. Here, we build on this work and show that it is possible to produce wt PV-3 and thermally-stabilised PV-3 (referred to as PV-3 SC8) VLPs in the native conformation (D Ag) using *Pichia pastoris*. We show that the PV-3 SC8 VLPs provide a much-improved D:C antigen ratio as compared to wt PV-3, whilst exhibiting greater thermostability than the current IPV vaccine. Finally, we determine the cryo-EM structure of the yeast-derived PV-3 SC8 VLPs and compare this to previously published PV-3 D Ag structures, highlighting the similarities between these recombinantly-expressed VLPs and the infectious virus, further emphasising their potential as a next-generation vaccine candidate for PV.

## Introduction

Poliovirus (PV) is the causative agent of poliomyelitis, an acute infectious disease which can result in paralysis and be fatal. However, since the establishment of the Global Polio Eradication Initiative (GPEI) in 1988 there has been >99% reduction in the number of paralytic poliomyelitis cases globally, with wild-type (wt) PV now only endemic in Afghanistan and Pakistan (1). The success of the GPEI is largely due to the use of two highly effective PV vaccines, the live-attenuated oral PV vaccine (OPV) and the inactivated PV vaccine (IPV) (2). Both of these vaccines target all three serotypes of PV (PV-1, PV-2 and PV-3), and although wt PV-2 and wt PV-3 were declared eradicated in 2015 and 2019, respectively (3), vaccine-derived viruses remain in circulation. Yet, despite the clear success of OPV and IPV, there are biosafety concerns surrounding the continued use of these vaccines as we move towards a ‘polio-free’ world, as both have the potential to re-introduce PV into the environment.

IPV provides excellent protection against poliomyelitis, however, as it does not induce sterilising immunity and cannot stop the virus from spreading within a population (4). Additionally, the production of IPV requires the cultivation of large amounts of infectious virus, which presents a considerable biosafety risk, as accidental release of such concentrated virus could have catastrophic results (5, 6). OPV has been instrumental in the near-eradication of PV, however, due to its inherent genetic instability, the attenuated virus can quickly revert to virulence. This reversion can cause vaccine-associated paralytic poliomyelitis (VAPP) which, in low vaccine coverage areas, can lead to circulating vaccine-derived PV (cVDPV) (7). Unfortunately, cVDPV cases now consistently outnumber wt PV cases worldwide (1). Furthermore, OPV can recombine with other polioviruses or polio-like enteroviruses during co-infection to generate novel neurovirulent chimeric viruses. This, coupled with reintroduction of infectious poliovirus into the environment via the chronic shedding of VDPV by immunocompromised individuals, highlights the risks associated with continued OPV usage as we strive towards eradication (8, 9).

PV is a 7.5 kb positive-strand RNA virus belonging to the *Enterovirus C* species. The PV genome contains two overlapping open-reading frames (ORFs), and whilst the function of the recently discovered uORF remains under investigation, it has been shown to be of importance in *ex vivo* human enteroid infection in the context of the prototypic *Enterovirus B* species member, Echovirus 7 (10). The major ORF is translated as a single polypeptide, containing 3 distinct regions, P1 (viral structural proteins), P2 and P3 (non-structural proteins required for proteolytic cleavage and viral replication). This polypeptide is then cleaved into mature viral proteins by the virally encoded proteases, 2A^pro^, 3C^pro^ and 3CD (11). The viral protease precursor protein, 3CD, has been shown to be primarily responsible for the cleavage of P1 into the individual capsid proteins, VP0, VP3 and VP1 (12, 13). Mature virions undergo a further cleavage event of VP0 into VP4 and VP2, which is associated with the encapsidation of viral RNA resulting in increased particle stability (14, 15).

Mature PV virions are comprised of 60 copies of VP1-VP4, which assemble as a ∼30 nm icosahedral capsid (16). The capsids incorporate a lipid molecule into a pocket within the VP1 protein, which is host-derived and important for stability (17). During PV infection, empty capsids (ECs), which do not contain viral genome and do not cleave VP0, are also produced. These particles are antigenically indistinguishable from mature viral particles (18). ECs have long been considered as potential virus-like particle (VLP) vaccines to replace the current PV vaccines, however, recombinantly-expressed wt ECs are inherently unstable and readily convert to an expanded form of the icosahedral particle (17, 18). This minor expansion has significant consequences for the antigenicity of ECs, converting these from the native antigenic form (termed D Ag) to the non-native form (termed C Ag). Unlike the D Ag form, the C Ag form is unable to induce a protective immune response to PV, therefore recombinant VLP vaccines against PV must retain the D Ag conformation (18-20).

VLPs mimic the structures of infectious virions but lack the viral genome, therefore making them safe and attractive options as recombinant vaccines, as evidenced by the licensed hepatitis B virus (HBV) and human papillomavirus (HPV) vaccines produced in yeast and yeast or insect cells, respectively (22–24). In the past 5 years, PV VLPs which maintain native antigenicity have been produced in several different systems, including mammalian, plant, and insect cells (25–30). Each of these systems can incur high development and production costs, making them less accessible in lower-to-middle income countries (LMICs) (31, 32). Yeast as a PV VLP production system has high potential for technology transfer to LMICs due to low costs of media. Furthermore, PV VLP production can utilise existing infrastructure, used to produce the HBV and HPV vaccines (33).

We have previously reported production of PV VLPs using *Pichia pastoris*, however, these VLPs were in the C Ag conformation (34). Here, we build on this work to demonstrate the production of wt PV-3 and stabilised PV-3 (referred to as PV-3 SC8), D antigenic VLPs in *Pichia pastoris*. We show that the PV-3 SC8 VLPs provide a much-improved D:C antigen ratio with greater thermostability than the current IPV vaccine. We also show that yeast-derived VLPs do not contain detectable levels of nucleic acid, therefore highlighting their potential as a safe alternative to the current PV vaccines. Finally, we determine the cryo-EM structure of the Yeast-derived PV-3 SC8 VLPs and compare it to previously published PV-3 D Ag structures (27, 30), highlighting the similarities between these recombinantly-expressed VLPs and the infectious virus.

## Methods

### Cells and viruses

HeLa cells were obtained from the National Institute of Biological Standards and Controls (NIBSC), UK. PichiaPink™ yeast strain one (Invitrogen, USA) was grown according to the instructions of the manufacturer. The infectious clone of wt PV-1 (Mahoney strain) used in this study was sourced from Bert Semler, University of California, USA. The cDNA was cloned downstream of a T7 RNA promoter to allow *in vitro* RNA synthesis. Additionally, a hammerhead ribozyme was included at the 5’ end allowing the production of an authentic infectious PV-1 RNA (35). To recover infectious virus, L-cells (which lack PV receptor, CD155) were transfected with PV-1 RNA and the resulting viruses were propagated in HeLa cells.

### Vector construction

The P1 gene of wt PV3 Saukett was amplified from a pT7RbzLeonSktP1_deletion mutant plasmid sourced from NIBSC, UK whereas PV3 SC8 P1 and an uncleavable 3CD gene were codon optimised for expression in *Pichia pastoris*. Both P1 genes and the uncleavable 3CD were cloned separately into the pPink-HC expression vector multiple cloning site (MCS) using *EcoRI* and *FseI* (NEB). Subsequently, the dual promoter expression vector was constructed through PCR amplification from position 1 of the 3CD pPink-HC to position 1285 inserting a *SacII* restriction site at both the 5’ and 3’ end of the product. The P1 expression plasmids were linearised by *SacII* (NEB), followed by the insertion of the 3CD PCR product into *SacII*-linearized P1 plasmid. All PCR steps were carried out with Phusion polymerase (NEB) using the manufacturer’s guidelines.

### Yeast transformation and induction

Plasmids were linearized by *Afl*II digestion (NEB) and then transformed into PichiaPink™ Strain one (Invitrogen, USA) by electroporation as per the manufacturer’s guidelines. Transformed yeast cells were plated on *Pichia* Adenine Dropout (PAD) selection plates and incubated at 28°C until sufficient numbers of white colonies appeared (3-5 days). To screen for high-expression clones, 8 colonies were randomly selected for small-scale (5 mL) expression experiments. Briefly, colonies were cultured in YPD for 48 hours at 28°C with shaking at 250 rpm, each culture was pelleted at 1500 × *g* and resuspended in YPM (1 mL & methanol 0.5% v/v) to induce protein expression and cultured for a further 48 hours. Cultures were fed methanol to 0.5% v/v 24 h post-induction. To determine expression levels of each clone the samples were analysed by immunoblotting as described below. For VLP production, a stab glycerol stock of a previously high-expressing clone was cultured for 48 hours in 5 mL YPD to high density. To increase biomass, 4 mL of the starter culture was added to 200 mL YPD in a 2 L baffled flask and cultured at 28°C at 250 rpm for a further 24 h. Cells were pelleted at 1500 × *g* and resuspended in 200 mL YPM (methanol 0.5% v/v) and cultured for a further 48 h. Cultures were fed methanol to 0.5% v/v 24 h post-induction. After 48 h cells were pelleted at 2000 × *g* and resuspended in breaking buffer (50 mM sodium phosphate, 5% glycerol, 1 mM EDTA, pH 7.4) and frozen prior to processing.

### Sample preparation and immunoblotting

Gradient fraction samples were prepared through a 1:5 mixture with 5x Laemmli buffer. Protein extracts were analysed by 12% SDS-PAGE (w/v) using standard protocols. Immunoblot analyses were performed using a monoclonal blend of primary antibodies against the VP1 protein of each PV1, PV2, and PV3 (Millipore MAB8655) followed by detection with a goat anti-mouse secondary antibody conjugated to horseradish peroxidase, and developed using the chemiluminescent substrate (Promega). To identify VP0, a rabbit polyclonal antibody (a kind gift from Ian Jones) was used followed by detection with a goat anti-rabbit secondary antibody conjugated to horseradish peroxidase, and developed using a chemiluminescent substrate (Promega) (36).

### Purification and concentration of PV and PV VLPs

Virus-infected HeLa cells were freeze-thawed and clarified by differential centrifugation. Supernatant was collected and virus pelleted through 30% (w/v) sucrose cushion at 151,000 × *g* (using a Beckman SW 32 Ti rotor) for 3.5 hours at 10°C. Virus pellet was resuspended in phosphate buffered saline (PBS) and clarified by differential centrifugation. Supernatant was purified through 15-45% (w/v) sucrose density gradient by ultracentrifugation at 151,000 × *g* (using a Beckman SW 40 rotor) for 3 hours at 10°C (18).

*P. pastoris* cell suspensions were thawed and subjected to cell lysis using CF-1 cell disruptor at ∼275 MPa chilled to 4°C following the addition of 0.1% Triton-X 100. The resulting lysate was centrifuged at 5000 rpm to remove the larger cell debris, followed by a 10,000 × *g* spin to remove further insoluble material. The resulting supernatant was nuclease treated using 25 U/mL DENARASE® (c-LEcta) for 1.5 hours at RT with gentle agitation. The supernatant was mixed with PEG 8000 (20% v/v) to a final concentration of 8% (v/v) and incubated at 4°C overnight. The precipitated protein was pelleted at 5,000 rpm and resuspended in PBS. The solution was pelleted again at 5,000 rpm and the supernatant collected for a subsequent 10,000 × *g* spin to remove any insoluble material. The clarified supernatant was collected and pelleted through a 30% (w/v) sucrose cushion at 151,000 × *g* (using a Beckman SW 32 Ti rotor) for 3.5 hours at 10°C. The resulting pellet was resuspended in PBS + NP-40 (1% v/v) + sodium deoxycholate (0.5% v/v) and clarified by centrifugation at 10,000 × *g*. The supernatant was collected and purified through 15-45% (w/v) sucrose density gradient by ultracentrifugation at 151,000 × *g* (using a 17 mL Beckman SW32.1 Ti rotor) for 3 hours at 10°C (18). Gradients were collected in 1 mL fractions from top to bottom and analysed for the presence of VLPs through immunoblotting and ELISA.

Peak gradient fractions as determined by immunoblotting and ELISA were then concentrated to ∼100 uL in PBS + 20 mM EDTA using 0.5 mL 100 kDa centrifugal concentration filters (Amicon) as per the manufacturer’s instructions.

### Thermostability Assay

The thermostability of VLPs was assessed using previously established assays (37). Briefly, previously quantified PV VLPs were diluted in Phosphate Buffered Saline (Corning 46-013-CM) to provide a uniform quantity of D antigen. Duplicate aliquots were incubated on ice (control) or in a thermocycler (BIO-RAD T100) at temperatures between 37 °C and 60 °C for 10 minutes.

Thermostability of the VLPs was assessed by measuring loss of D Ag by ELISA, detection of D antigenic particles was determined through PV-3 specific Mab 520.

### Enzyme-linked immunosorbent assay (ELISA)

To determine antigen level in gradient fractions a non-competitive sandwich ELISA was used to measure PV3 D and C antigen content (38). Briefly, two-fold dilutions of antigen were captured using a PV3-specific polyclonal antibody, and detected using PV3-specific, D antigen (Mab 1050) or C antigen (Mab 517.3) specific monoclonal antibodies (kindly provided by NIBSC), followed by anti-mouse peroxidase conjugate (39, 40). To determine the presence of individual antigen sites concentrated samples were captured using a PV3-specific polyclonal antibody and detected using antigen site-specific monoclonal antibodies, kindly provided by NIBSC, followed by anti-mouse peroxidase conjugate. The specific antigen sites are listed in table 1. BRP (Sigma) was used as the standard for D antigen content in each ELISA. All ELISAs were analysed through Biotek PowerWave XS2 plate reader.

**Table 1.**
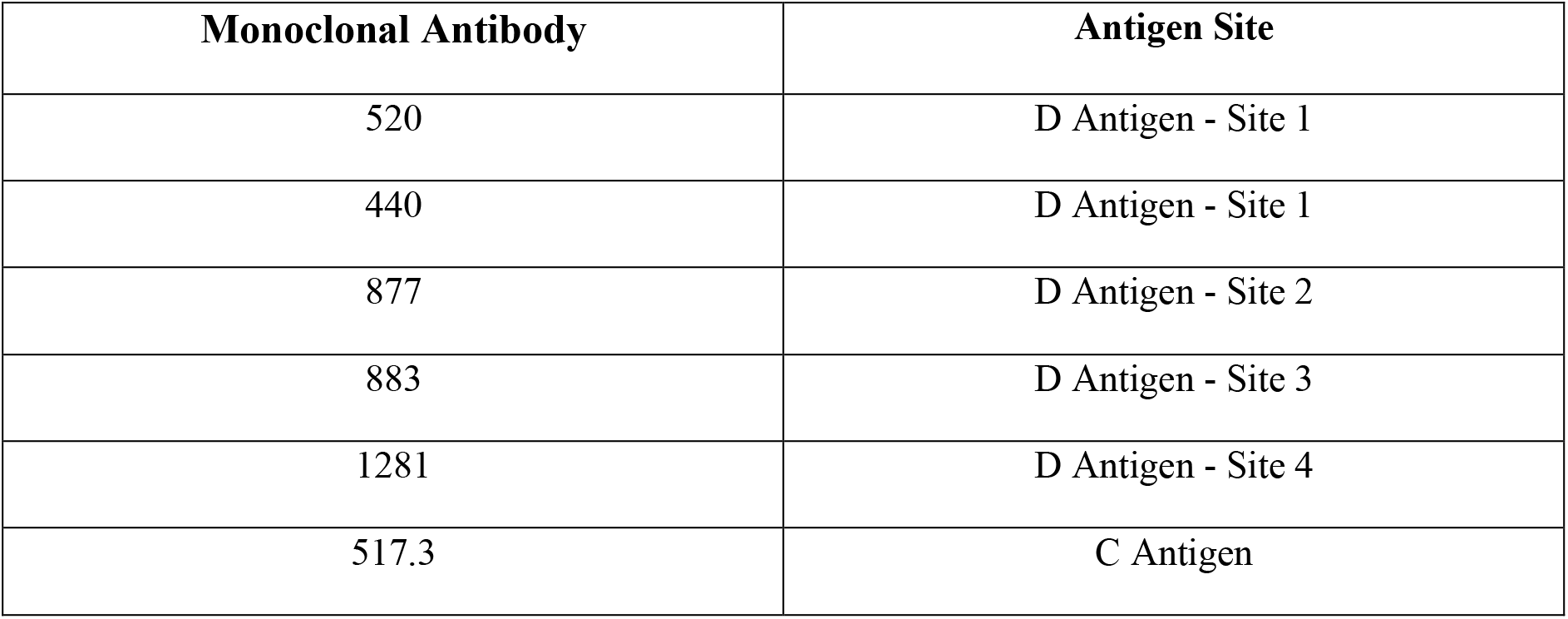
List of monoclonal antibodies and corresponding site recognition information (38, 40).

### Particle Stability Thermal Release Assay (PaSTRy)

PaSTRy assays (41) were used to compare the presence of RNA within the purified PV-3 VLPs with infectious PV using the nucleic acid binding dye SYTO9. 500 µg of purified wt PV-3 VLPs, PV-3 SC8 VLPs or infectious PV-1 were incubated with 5 µM SYTO9 (Thermo Fisher) and SYPRO red (Thermo Fisher) at a 6x final concentration in PBS. The incubation was carried out on a temperature ramp from 25°C to 95°C and the fluorescent signal was measured at 1°C intervals every 30 seconds using the Stratagene MX3005P qPCR machine (Aligent Technologies).

### Negative stain electron microscopy

To prepare samples for negative stain transmission EM, carbon-coated 300-mesh copper grids were glow-discharged in air at 10 mA for 30 seconds. 3 µl aliquots of purified VLP stocks were applied to the grids for 30 seconds, then excess liquid was removed by blotting. Grids were washed twice with 10 µl distilled H_2_O. Grids were stained with 10 µl 1% uranyl acetate solution, which was promptly removed by blotting before another application of 10 µl 1% uranyl acetate solution for 30 seconds. Grids were subsequently blotted to leave a thin film of stain, then air-dried. EM was performed using an FEI Tecnai G2-Spirit transmission electron microscope (operating at 120 kV with a field emission gun) with a Gatan Ultra Scan 4000 CCD camera (ABSL, University of Leeds).

### Negative stain image processing

Raw micrographs were visualised with ImageJ 1.51d (42, 43).

### Cryo-EM sample preparation and data collection

Sucrose gradient purified fractions of yeast-derived PV-3 SC8 were pooled and buffer exchanged into PBS + 20mM EDTA (pH 7) using Zeba Spin Desalting Columns with a 7K molecular weight cut-off (MWCO) (Thermo Fisher Scientific) and concentrated using Amicon Ultra centrifugal filter devices (100 kDa MWCO, Merck Millipore) to a final concentration of ∼0.5 mg/ml. Three to four microliters of PV-3 SC8 VLP was applied to glow-discharged Lacey carbon copper grids with an ultra-thin carbon support film (Agar Scientific). After 30 s unbound sample was removed by manual blotting with filter paper. To increase the number of particles in the holes, grids were re-incubated with a further 3-4 μl of sample for 30 s, followed by mechanical blotting for 4 s and rapid vitrification in liquid ethane with a Vitrobot Mark IV plunge-freezing device (Thermo Fisher Scientific) operated at 4 °C and 100 % relative humidity.

Cryo-EM data acquisition was performed at 300 kV with a Titan Krios G3i microscope (Thermo Fisher Scientific) at the OPIC electron microscopy facility, UK. The microscope was equipped with a K2 Summit (Gatan) direct electron detector (DED) and an energy filter (GIF Quantum, Gatan) operating in zero-loss mode (0-20 eV energy selecting slit width). Micrographs were collected as movies using a defocus range of -2.4 μm to -0.9 μm in single-electron counting mode with a pixel sampling of 0.82 Å per pixel resulting in a calibrated magnification of ×60,975. Data were collected using SerialEM (44). Data acquisition parameters are summarized in Supplementary Table 1.

### Cryo-EM image processing

Image processing and single-particle reconstruction was performed using RELION-3.1 (45) unless indicated otherwise. Individual movie frames were aligned and averaged with dose weighting using MotionCor2 (46) to produce images compensated for electron beam-induced specimen drift. Contrast transfer function (CTF) parameters were estimated using CTFFIND4 (47) Micrographs showing astigmatism or significant drift were discarded. Particle-picking was performed using crYOLO (48) by first training the neural network on a randomly selected subset of ∼100 manually picked particles from 50 micrographs covering a range of defocus values. Once trained crYOLO was used to pick the complete dataset in an automated manner, and the saved particle coordinates were then imported into RELION.

Single-particle structure determination used established protocols in RELION for image classification and gold-standard refinement to prevent over-fitting (49) Picked particles (numbers given in Supplementary Table 1) were subjected to two rounds of reference-free two-dimensional classification to discard bad particles and remove junk classes. The particle population was further enriched by three-dimensional (3D) classification to remove broken and overlapping particles. The starting reference model was the previously determined cryo-EM structure of PV-3 SC8 from a plant expression system (30) (EMDB accession code EMD-3747) low-pass filtered to 60 Å to avoid bias.

A final set of particles (numbers given in Supplementary Table 1) were selected from the best aligned 3D class averages for high-resolution 3D auto-refinement with the application of icosahedral symmetry throughout. A representative class from the end of 3D classification was low pass filtered to 40 Å to avoid bias and used as a reference during refinement. After the first round of refinement the data were subjected to CTF refinement to estimate beam tilt, anisotropic magnification, per-particle defocus and astigmatism, and also Bayesian polishing of beam-induced motion-correction with default parameters (45). This procedure was performed twice with 3D auto-refinement after each round. The final resolution was estimated using a Fourier shell correlation (FSC) threshold of 0.143 (50). The cryo-EM maps were sharpened using Post-processing in RELION by applying an inverse B-factor of -59.1 Å^2^. Local resolution was estimated using the RELION implementation of local resolution algorithm (51) and locally scaled maps were used for model building and refinement in all cases. Data processing statistics are summarized in Supplementary Table 1.

### Atomic model building, refinement and analysis

The atomic coordinates of the previously determined structure of PV-3 SC8 (PDB 6Z6W) were manually placed into the cryo-EM electron potential map using UCSF Chimera (52). Manual fitting was optimised with the UCSF Chimera ‘Fit in Map’ command (52). Manual rebuilding was performed using the tools in Coot (53), followed by iterative positional and B-factor refinement in real-space using phenix.real_space_refine (54) within Phenix (55). All refinement steps were performed in the presence of hydrogen atoms and only the atomic coordinates were refined; the map was kept constant. Each round of model optimization was guided by cross-correlation between the map and the model. The final model was validated using the MolProbity (56) tools within Phenix (55). Refinement statistics are shown in Supplementary Table 2. Structural superpositions of PV-3 capsid protomers were performed with program SHP (57) and molecular graphics were generated using Pymol (The PyMOL Molecular Graphics System, Version 2.0 Schrödinger, LLC.) and UCSF ChimeraX (58).

### Statistical Analysis

All t-tests were two-tailed and performed using the statistical analysis software Prism.

## Results

### Production of *wt* and stabilised PV3 VLPs using *Pichia pastoris*

Production of enterovirus VLPs such as Coxsackie A6, Coxsackie A16 and enterovirus A71 by recombinant expression has been demonstrated previously (59–61). Furthermore, we have previously reported the production of PV-1 VLPs in *Pichia pastoris* using a variety of expression cassettes. However, these VLPs were in the non-native C antigenic conformation and therefore incapable of inducing protective immune responses (34). Here, we demonstrate the production of D antigenic PV-3 VLPs in *P. pastoris*. We employed a dual promoter expression system for the structural precursor protein (P1) and the viral protease, 3CD, which cleaves P1 into VP0, VP3 and VP1, therefore allowing for capsid assembly (Fig. 1B). This system was employed to produce wt PV-3 Saukett (Skt) and PV-3 Skt SC8 VLPs (Herein, referred to as wt PV-3 and PV-3 SC8 respectively). The PV-3 capsid mutant SC8 has previously been shown to ‘lock’ empty capsids in the D Ag conformation when grown in mammalian tissue culture as an infectious clone and has since been shown to maintain this conformation in recombinant VLPs produced using plant and modified vaccinia Ankara (MVA)-based mammalian expression systems (27, 30, 37). Therefore, we determined the D antigenic characteristics of PV-3 SC8 VLPs produced in *P. pastoris* as a highly-tractable heterologous expression system.

**Figure 1:**
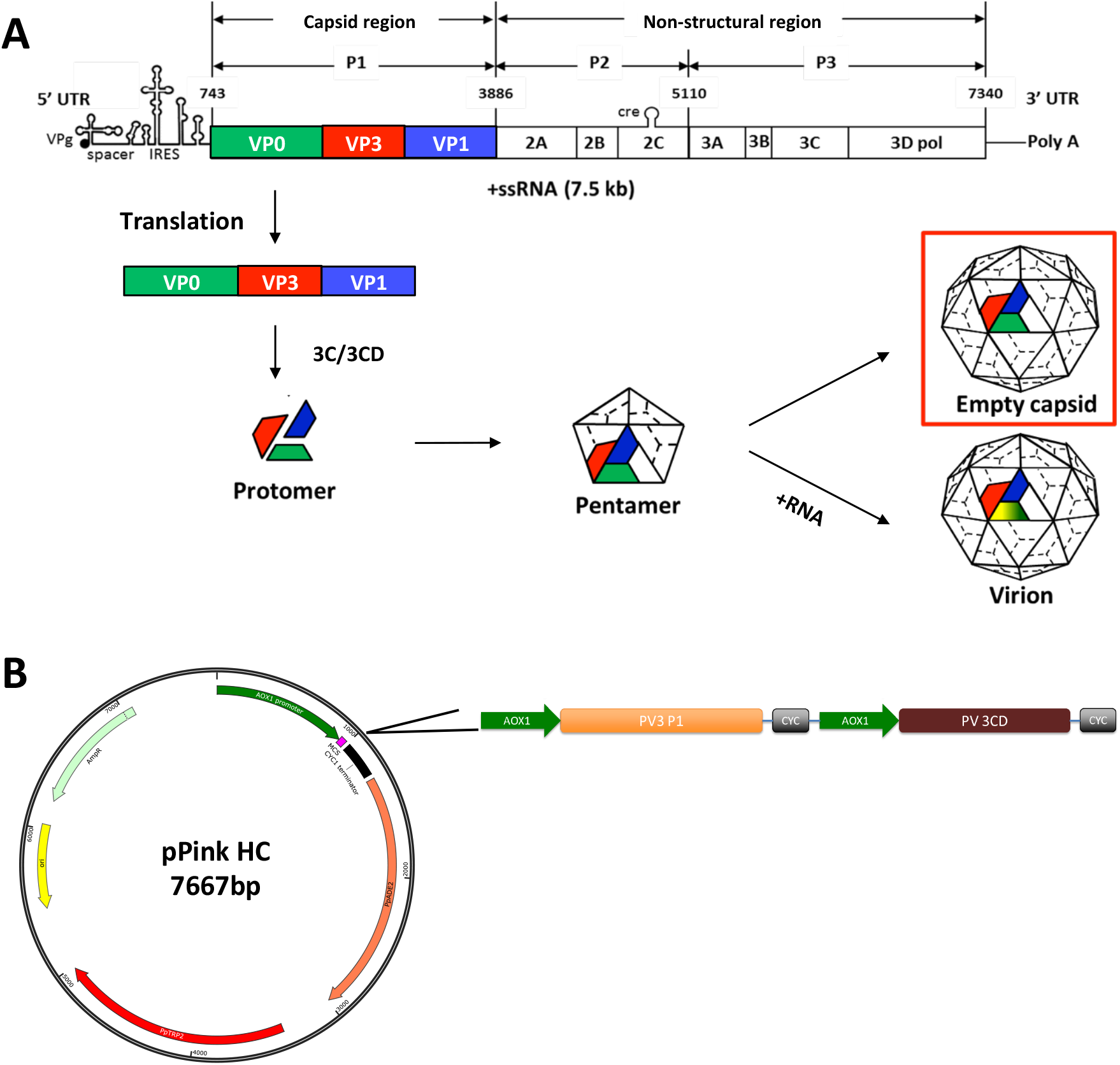
Schematic of poliovirus genome and expression cassette. **A**: Schematic highlighting the structural and non-structural regions of the PV genome. The capsid region P1 is cleaved post-translationally by the viral protease (3CD) to produce VP0 (green), VP3 (red), and VP1 (blue), which form protomers, followed by pentamer formation, which culminates in the production of either virions in the presence of viral RNA or empty capsids, in which VP0 remains uncleaved. **B**: The pPink HC expression vector with the dual alcohol oxidase (AOX) promoter expression construct, which drives the expression of the P1 capsid protein and the viral protease, 3CD.

High-expressing *P. pastoris* clones for both wt PV-3 and PV-3 SC8 were cultured in 2x 200 mL culture volumes, induced with 0.5% methanol (v/v) and the cell pellets collected 48 hours (h) post-induction. To determine the levels of VLP production by each construct, cell pellets were homogenised at ∼275 MPa and the resultant lysates purified through multiple rounds of centrifugation culminating in 15-45% sucrose gradients. Following ultracentrifugation, gradients were fractionated and assessed by immunoblot for the presence of viral capsid proteins VP1 and VP0 (Fig. 2a). The VLPs migrated to the middle of the gradients (peak between fractions 8 and 10), as detected by both anti-VP1 and anti-VP0 western blotting. Despite similar culture volumes and cell pellet weights, wt PV-3 VLPs were expressed at higher levels than PV-3 SC8 VLPs, suggesting that the stabilising mutations present in PV-3 SC8 resulted in reduced translation or efficiency of particle assembly.

**Figure 2:**
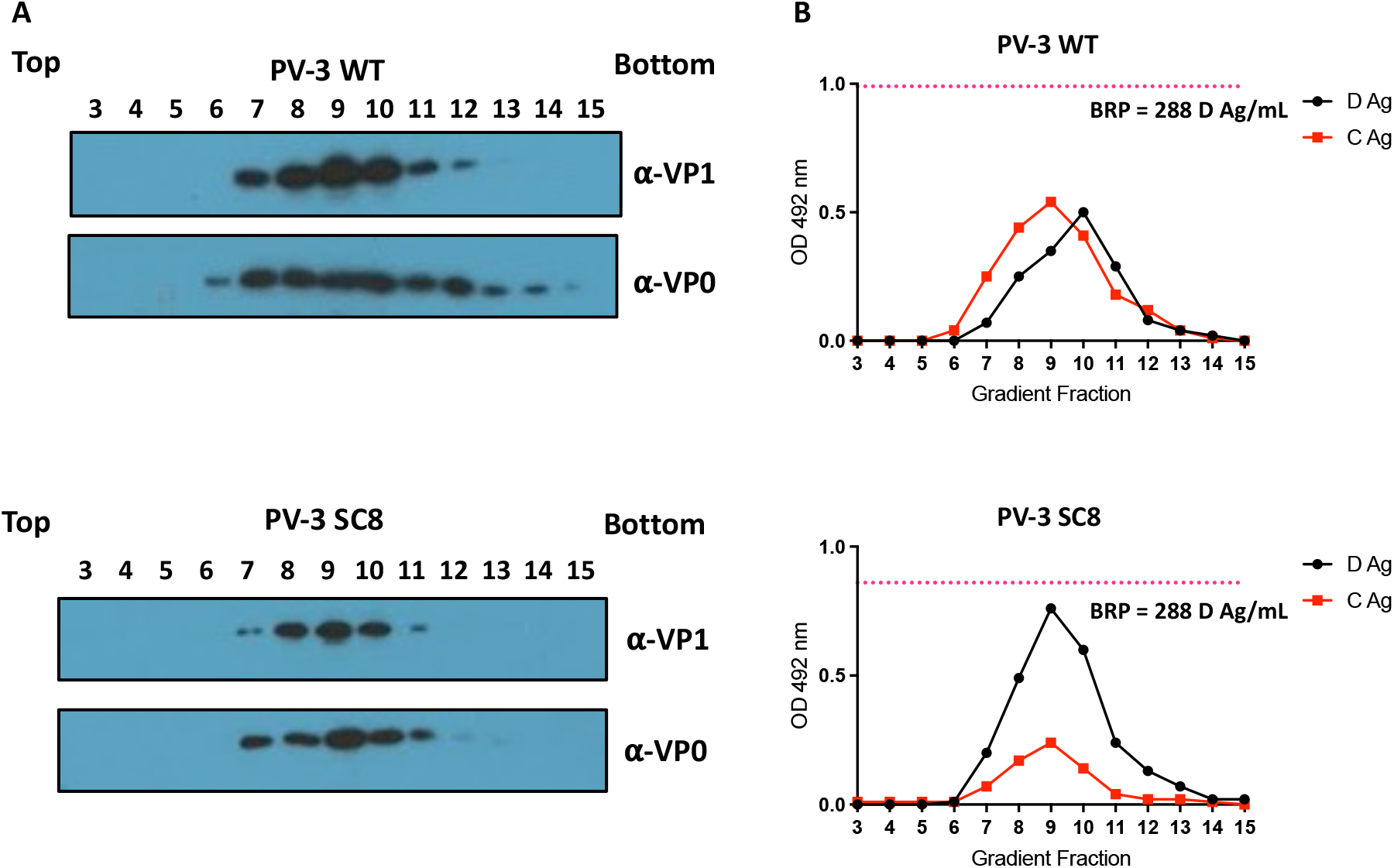
Purification and antigenicity of PV-3 WT and PV-3 SC8 VLPs. **A:** All samples were purified by ultracentrifugation and fractioned from top to bottom in 1 mL aliquots. A 12 µL sample from each fraction was then taken and boiled in 5x Laemmli buffer and separated by SDS PAGE prior to analysis by immunoblot using mouse monoclonal α-VP1 (Millipore MAB8655) and rabbit polyclonal α-VP1. **B**: Antigenicity of PV-3 WT and PV-3 SC8 VLPs. Reactivity of gradient fractions with Mab 520 (D antigen) and Mab 517.3 (C antigen) in ELISA. The pink dashed line represents the positive control, BRP, for the D antigen ELISA. OD 492 nm is represented in arbitrary units. The figure is a representative example of three separate experiments for each construct.

Gradient fractions were then assessed for antigenic characteristics by enzyme-linked immunosorbent assay (ELISA) (Fig. 2B). Fractions were analysed using a standard protocol, as established by the National Institute for Biological Standards and Control (NIBSC), using the current inactivated vaccine standard (BRP) as a positive control. wt PV-3 and PV-3 SC8 VLPs included both D and C antigenic particles and the peak antigen content determined by ELISA in Fig. 2B, mirrored the peaks detected by immunoblot in Fig. 2A. However, the ratio of D:C Ag was strikingly different, with wt PV-3 VLPs at a ∼1:1 ratio whereas PV-3 SC8 VLPs were at ∼2.5:1 ratio, highlighting the impact of the stabilising mutations on the ability of PV-3 VLPs to maintain the D Ag conformation.

### PV-3 SC8 VLPs are thermally stable with antigenic profiles indistinguishable from current vaccine

As both wt PV-3 and PV-3 SC8 included D Ag VLPs, we determined their antigenic similarity to the current inactivated vaccine, using the BRP standard described above. We employed several monoclonal antibodies (Mabs) to determine the presence or absence of the major antigenic sites (described in Table 1) on wt PV-3 and PV-3 SC8 VLPs in comparison to BRP (Fig. 3A) (38, 40). The wt PV-3 VLPs showed reduced level of D Ag reactivity compared to BRP and PV-3 SC8 with each D Ag Mab tested, aside from Mab 1281, although these differences were not significantly different. Additionally, the wt PV-3 VLPs reacted significantly more strongly with a C Ag specific Mab compared to BRP (*p* = 0.0370). The PV-3 SC8 VLPs had D Ag levels similar to those seen with BRP across the antigenic sites. Although PV-3 SC8 VLPs did show some increased C Ag reactivity in comparison to BRP, this difference was not statistically significant. Interestingly, both wt PV-3 and PV-3 SC8 showed increased reactivity to Mab 440 in comparison to BRP, although this difference was only significant for PV-3 SC8 (*p* = 0.0224). This was expected, as the process of inactivation employed during BRP production destroys the antigenic site recognised by Mab440 and is therefore referred to as a Sabin-specific antigenic site (62). Taken together, these data suggests that the PV-3 SC8 VLPs are comparable in their antigenicity to the current vaccine and may offer the potential of increased immunogenicity through the preservation of the Sabin-specific antigenic site 1.

**Figure 3:**
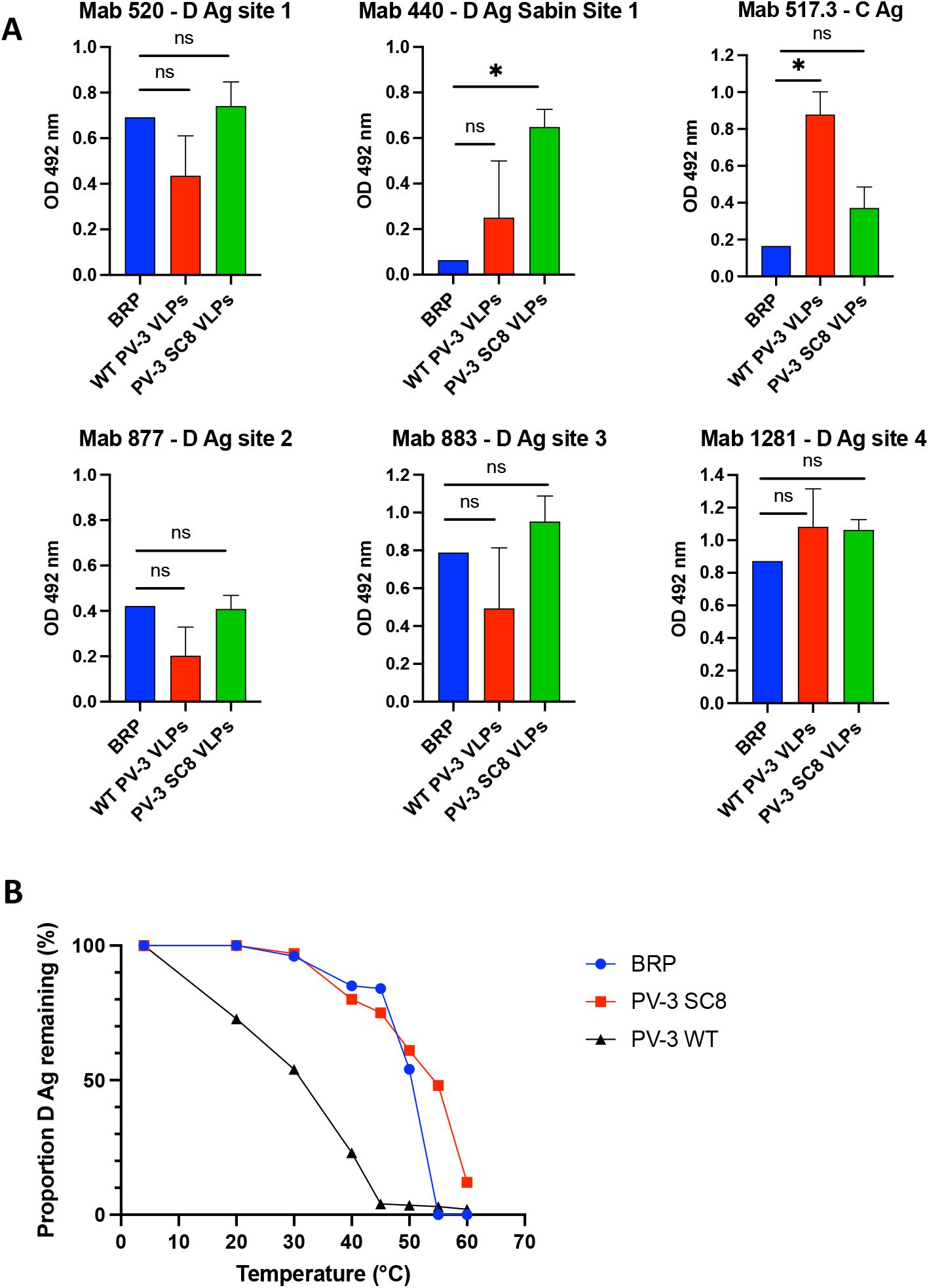
Antigen mapping and thermostability. **A**: Antigen mapping of *yeast-derived* PV-3 wt and PV-3 SC8 VLPs. Reactivity of concentrated VLPs and BRP with PV3 Mabs specific for different antigenic sites by ELISA. OD 492 nm is represented in arbitrary units (n=3) Means +/-standard deviation. Statistical analysis determined by two-tailed T-test (ns, not significant; * P-value > 0.05) **B**: Reactivity of PV-3 SC8 VLP, PV-3 wt VLPs and BRP aliquots to D-antigen specific Mab 520 in ELISA after incubation at different temperatures for 10 minutes, normalised to corresponding aliquot incubated at 4 °C. The figure is a representative example of three separate experiments.

Next, we determined the thermostability of yeast-derived VLPs in comparison to BRP as potential next-generation vaccines will need to match, or preferably surpass the thermostability of the current vaccine in order to address the issues surrounding the maintenance of cold-chain storage for vaccine administration within LMICs. Therefore, wt PV-3 VLPs, the stabilised PV-3 SC8 VLPs and BRP were subjected to increasing temperatures and assessed for the retention of D antigenicity (Fig. 3B). As expected, wt PV-3 VLPs were less thermostable than both PV-3 SC8 VLPs and BRP, with only 23% D Ag remaining at ∼40°C. PV-3 SC8 VLPs showed increased thermostability in comparison to BRP, with BRP showing a 50% decrease in D antigenicity at ∼51°C compared to ∼53°C for PV-3 SC8 VLPs, which still maintained 48% D Ag at 55°C, at which temperature BRP no longer contained detectable D Ag.

### Yeast-derived VLPs maintain icosahedral morphology and do not package detectable levels of nucleic acid

Figure 4A shows representative micrographs of yeast-derived wt PV-3 VLPs and PV-3 SC8 VLPs. The negative-stain EM images demonstrate that the VLPs maintain the classical icosahedral morphology associated with picornaviral particles, measuring ∼30 nm in diameter, consistent with previous EM images of infectious poliovirus virions (34). Both sets of VLP micrographs show some smaller particles, these impurities are likely to be alcohol oxidase or fatty acid synthetase which have been shown to co-purify with VLPs through the sucrose gradients (63).

**Figure 4:**
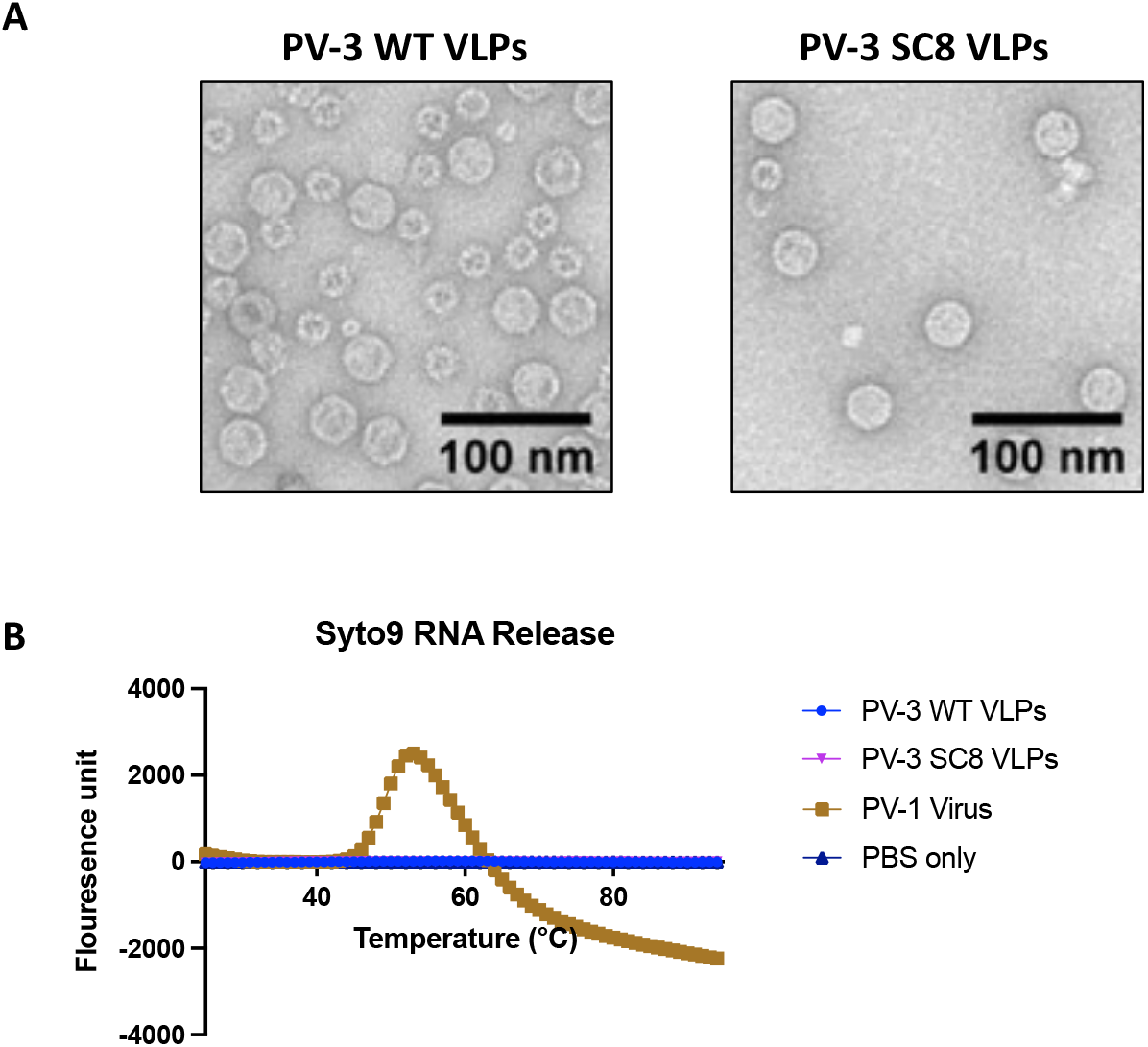
Negative stain transmission electron microscopy of yeast-derived VLPs and PaSTRy. **A:** Representative micrographs from preparations of PV-3 wt and PV-3 SC8 VLPs, stained with uranyl acetate. Scale bar shows 100 nm. **B**: Particle Stability Thermal Release assay (PaSTRy) equal amounts of either VLP or infectious virus samples were assessed for RNA release over an increasing temperature gradient. PBS was used as a negative control and fluorescence is represented in arbitrary units. The figure is a representative example of three separate experiments.

VLP vaccines lack viral genetic information, and hence are replication-incompetent. Therefore, we determined whether the yeast-derived PV VLPs had packaged substantial levels of either cognate mRNA or any non-specific yeast RNA. Concentrated VLPs were assessed for the presence of nucleic acid and compared to infectious PV-1 using a particle stability thermal release assay (PaSTRy, (41)) (Fig. 4B). As VLPs or viruses are heated, they undergo conformational changes which expose the internalised nucleic acid(s). Syto9 dye binds any exposed or released nucleic acid leading to the emission of a fluorescent signal. Fig. 4B shows that for the positive control, infectious PV-1 virions, there is a clear peak for RNA release at 54°C. However, there is no evidence of nucleic acid exposure or release from either the wt PV-3 VLPs or the PV-3 SC8 VLPs. This suggests that the VLPs produced in *P. pastoris* are reminiscent of ECs produced during natural virus infection and contain no nucleic acid.

### Comparative analysis of PV-3 SC8 Cryo-EM structures

The structure of yeast-derived PV-3 SC8 VLP was determined by cryo-EM to facilitate comparison with the same VLP produced previously in plant and mammalian expression systems (27, 30) and wild PV-3 virions. Single-particle reconstruction of a final set of 2712 particles from 3717 micrographs resulted in a 2.7 Å icosahedral reconstruction as assessed with the FSC 0.143 threshold criterion (50) (Supplementary Figure 1 and Supplementary Table 1). The cryo-EM map revealed well-ordered density for the VP0, VP1 and VP3 subunits of the capsid protein, allowing the bulk of the backbone and sidechain to be modelled and refined (Supplementary Figure 2). Comparison of the yeast-derived PV-3 SC8 structure against PV-3 SC8 from plant and mammalian cells showed that these particles have identical morphology. The diameter of the capsids (∼300A) corresponds to a D Ag, unexpanded particle, demonstrating that recombinantly produced PV-3 SC8 reliably assemble into the native antigenic state like natural PV virions. The PV-3 VLPs display the architectural features associated with native PV virions such as the ‘canyon’ depression around the fivefold axis and protrusions above the VP1, VP0 and VP3 subunits (Fig. 5A) (16). Native conformation (D Ag) PVs normally contain a lipid molecule in the hydrophobic pocket of the capsid protein VP1 subunit (17). For the yeast and mammalian PV-3 SC8 VLPs cryo-EM density was observed within the hydrophobic pocket of the VP1 β-barrel (Fig. 5B) corresponding to a host-derived lipid ‘pocket-factor’ and modelled as sphingosine (18 carbon length) as for the wt virus (17). However, differences were observed in the pocket factor density between yeast or mammalian expressed VLPs and the plant expressed PV-3 SC8 VLP where no cryo-EM density for a bound pocket factor could be observed (Fig. 5B) (30) despite the pocket being in an ‘open’ conformation as expected for a D Ag particle. This was attributed to the cryo-EM data reflecting low occupancy of a mixture of lipids from the plant cells (30).

**Figure 5:**
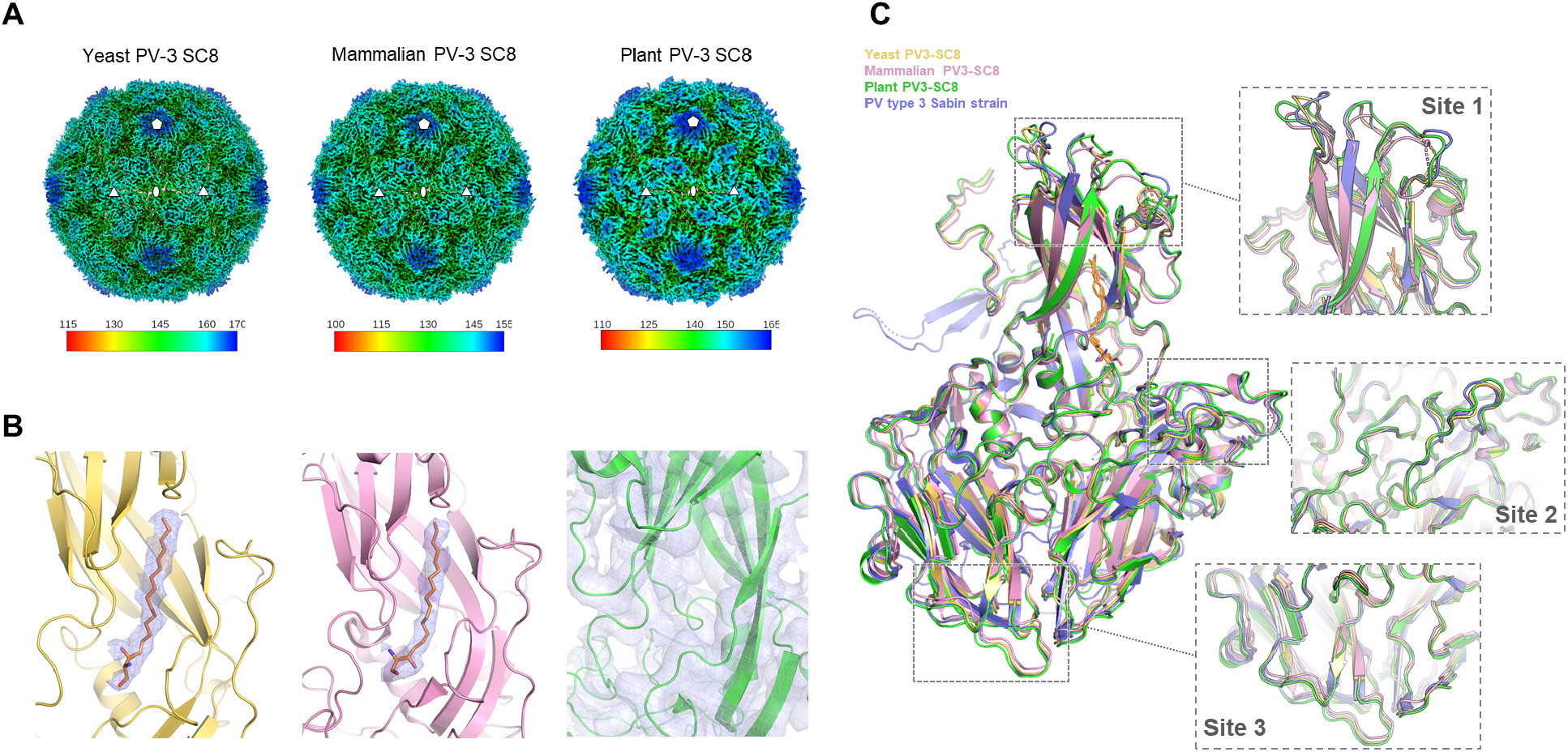
Comparison of PV-3 SC8 VLPs produced in plant, mammalian and yeast cells. A: Three dimensional cryo-EM reconstructions of PV-3 SC8 VLPs expressed in yeast, mammalian and plant cells. The VLPs are viewed along the icosahedral twofold symmetry axis and coloured by radial distance in Å from the particle center according to the colour keys provided. Five, three and twofold symmetry axes are labelled with symbols. B: Close-up views of the capsid protomer VP1 subunit hydrophobic pocket for the yeast (pale yellow), mammalian (pink) and plant (green) produced VLPs. Cryo-EM density maps are shown rendered around the VP1 protein for plant PV-3 SC8, or around the modelled sphingosine lipid in the yeast and mammalian PV-3 SC8 VLPs. C: Structural superposition of the capsid protomers of yeast (pale yellow, PDB 8ANW), mammalian (pink, PDB 6Z6W) and plant (green, PDB 5O5B) produced PV-3 SC8 VLPs with the structure of the PV-3 virus sabin strain (blue, PDB 1PVC). The sphingosine lipid modelled in the VP1 pockets of yeast and mammalian PV-3 SC8 VLPs and PV-3 sabin virus are shown as orange sticks. Boxes highlight antigenic sites on the VP1, VP2 and VP3 subunits of the capsid protein and are shown in expanded views labelled as site 1, site 2 and site 3 respectively.

The surface loops and C-termini of the capsid proteins which confer the antigenic characteristics are in identical conformations to the native PV3 virus structure (17), with root-mean-square deviations in Cα atoms of 0.93 Å, 0.99 Å and 0.48 Å between native PV3 virus and plant, mammalian and yeast derived PV-3 SC8 VLPs respectively (Fig. 5C). As expected the internal features resemble those of an empty capsid e.g. PDB:1POV (18) with disordered N-termini for VP1 (65 residues) and VP0 (including the entirety of what would be VP4 after cleavage). As observed for the PV-3 SC8 VLP produced in plant and mammalian cells, the capsid stabilizing mutations can be clearly identified in seven of the eight positions, the other residing in disordered VP4 (27, 30).

## Discussion

Both the current PV vaccines, OPV and IPV, require the production of substantial quantities of infectious virus, which carries significant biosafety risks due to the potential to reintroduce the virus to the environment (2, 4). Additionally, the use of OPV has contributed to the global increase in cVDPV cases, which now surpass those caused by wt PV (9). However, these vaccines are incredibly important in the effort to eradicate PV, especially with the recent licensing of nOPV2. This is a novel and intelligently designed vaccine which significantly reduces the likelihood of reversion or recombination, two of the major concerns surrounding the current OPV (64). With this newly available vaccine, there is a renewed hope that we are edging closer towards a polio-free world, and therefore there is a need for an alternative PV vaccine to maintain eradication by ensuring against accidental reintroduction of the virus. One potential solution is the production of virus-free VLP vaccines using a heterologous expression system. This approach has been shown to be successful in plant, insect and MVA-based mammalian expression systems, however, each of these requires specialist equipment, which in turn leads to increased production costs, making these systems difficult to transfer to LMICs (25–31). In contrast, yeast is a tractable heterologous expression system which can produce foreign proteins to similar levels as mammalian cells but at a fraction of the cost. As HBV and HPV vaccines are already being produced in LMICs, technology transfer of yeast-produced PV VLPs should be achieveable with affordable production costs (22, 23, 33). We have previously reported the production of PV-1 VLPs using *Pichia pastoris*, however, these particles were not in the correct D Ag conformation necessary to induce protective immunity against PV (34). Here, we demonstrate the production of D Ag PV-3 genetically stabilised VLPs using *P. pastoris*.

The wt PV-3 VLPs described here exhibited both C and D antigenicity (Fig. 2B). This is in contrast to the PV-1 VLPs described previously (34). The higher antigenic stability of wt PV-3 ECs as reported by Fox et al (37) likely explains this. For the wt PV-3 VLPs described here, the D:C ratio was 1:1, whereas with the stabilised PV-3 SC8 VLPs, this ratio was ∼2.5:1, showing that PV-3 SC8 includes a greater proportion of D Ag. However, we saw a reduced final yield of VLPs in the gradients by both immunoblot and ELISA (Fig. 2A) when using the PV-3 SC8 construct in comparison to the wt PV-3, as had also been previously observed when expressing these sequences in insect cells (29). This suggests that the PV-3 SC8 P1 sequence may impact mRNA translation or, following processing by 3CD, the stabilising mutations present within PV-3 SC8 may reduce the efficiency of capsid assembly. Fortunately, this reduction did not seem to negatively impact the overall amount of D Ag produced.

Following the observation that both wt PV-3 and PV-3 SC8 VLPs were produced in the native confirmation, we compared the antigen site maps of these yeast-derived VLPs with current inactivated vaccine (Fig. 3A). Interestingly, both wt PV-3 and PV-3 SC8 VLPs were reactive to Mabs detecting each of the major antigen sites, although PV3 SC8 VLPs had greater reactivity to D Ag Mabs than wt PV-3 VLPs. This, coupled with the reactivity to the C-specific Mab 517.3, highlights the importance of the stabilising mutations for producing D antigenic particles with reduced amounts of C Ag. Further to this, the yeast-derived VLPs were both reactive to Mab 440, which recognises the Sabin-specific antigenic site 1, whereas BRP was not reactive (as the inactivation process abolishes this site). Therefore, this suggests that PV-3 SC8 VLPs may provide an increased breadth of neutralising antibodies over the current IPV vaccine.

Any next-generation PV vaccine will need to be thermally stable to address the problems of storage and the challenges of maintaining a cold-chain when delivering the vaccine to remote places within LMICs. To this end, we compared the thermostability of the stabilised yeast-derived VLPs with BRP. PV-3 SC8 VLPs retained 50% D Ag to a higher temperature than the inactivated vaccine (53°C vs 51°C), and also retained 48% D Ag at 55°C whereas D Ag was no longer detectable at this temperature for BRP. The increased thermostability for the yeast-derived PV-3 SC8 VLPs was consistent with that previously published for PV-3 SC8 VLPs produced in mammalian and plant expression systems (27, 30). Interestingly, plant-and-yeast-derived PV-3 SC8 VLPs showed similar stabilities, losing 50% D Ag at 50°C and 53°C respectively, whereas the mammalian-produced PV-3 SC8 VLPs maintained 50% D Ag at temperatures above 60°C, suggesting that thermostability may also be influenced by the expression system in addition to the presence of stabilising mutations. However, as each expression system showed significantly increased thermostability compared to wt PV-3 empty capsids, we believe that these data provide further evidence that PV-3 SC8 VLPs have the potential to become part of a next-generation PV vaccine.

An important safety implication of VLP vaccines is that they do not package the viral genome, and therefore are not infectious. Consequently, it was of importance to determine if these VLPs contained cognate mRNA or host RNAs as this may have implications for regulatory approval. The PaSTRy assay highlighted that although a clear peak was seen for viral RNA release from infectious PV-1, there was no detectable nucleic acid released from either the wt PV-3 VLPs or the PV-3 SC8 VLPs, in line with our previous work which also suggested negligible levels of nucleic acids in Yeast-produced PV VLPs (34). Further work is required to determine the immunogenicity of these yeast-derived particles *in vivo* and to generate similarly stabilised PV-1 and PV-2 VLPs.

Structural data for PV-3 SC8 VLPs has now been generated for particles produced recombinantly in plants, mammalian and yeast cells (27, 30, 65) and confirms that PV-3 SC8 VLPs assemble into particles that adopt a native conformation akin to mature wt PV virions. VLPs from yeast and mammalian systems package a lipid into the VP1 β-barrel as in PV3 virions (17) but ordered cryo-EM density for these molecules was not observed in plant-made PV-3 SC8 VLPs (30). However, the pocket may be partially occupied by a mixture of lipids from plant cells since the pocket was not collapsed, indeed it was shown that synthetic ‘pocket factors’ could be bound in the VP1 pocket of plant-made PV-3 SC8 VLP (30). This observation has recently been extended to the PV-3 SC8 VLPs made in *P*. pastoris with ‘pocket factor’ compounds in the VP1 hydrophobic pocket and other potential stabilising additives at additional site (65). These results offer future strategies for additional methods to further stabilise recombinant PV VLP vaccine candidates.

In conclusion, we have shown that *P. pastoris* is a viable expression system for the production of D antigenic PV VLPs, which not only share a number of characteristics with the current vaccine but potentially improve upon it. Additionally, our data corroborates the results observed in other expression systems using this thermostable mutant, with the structural data highlighting the great similarities, and subtle differences, in particles produced from the different heterologous expression systems (27, 29, 30, 37). Overall, we believe that *P. pastoris* has the potential to produce VLP vaccines not only for a polio-free world but also as a model system for other members of the picornavirus family.

## Authors and Contributions

LS, DJR & NJS conceived and designed the experiments. LS, KG, JJS & MWB conducted the experiments. LS, KG, JJS, MWB, CP, EEF, DIS, DJR & NJS analysed the data. LS, DJR & NJS wrote the initial manuscript. KG, JJS, MWB, EEF & DIS reviewed and edited the manuscript. Funding was secured for this research by EEF, DIS, DJR and NJS.

## Data deposition

The model for PV-3 SC8 and cryo-EM map are deposited with PDB and EMDB accession codes: 8ANW and EMD-15543 respectively.

## Acknowledgments

We thank other members of the Stonehouse/Rowlands group, at the University of Leeds, for their insightful contributions. We also thank Bert Semler (University of California, USA) for the cDNA of wt PV-1 (strain Mahoney). This work was performed as part of a WHO-funded collaborative project towards the production of a virus-free polio vaccine involving the following institutions: University of Leeds, University of Oxford, University of Reading, University of Florida, Harvard University, John Innes Centre, the Pirbright Institute and the National Institute for Biological Standards and Control.

Computation used the Oxford Biomedical Research Computing (BMRC) facility, a joint development between the Wellcome Centre for Human Genetics and the Big Data Institute supported by Health Data Research UK and the National Institute for Health (NIHR) Oxford Biomedical Research Centre. Financial support was provided by the Wellcome Trust Core Award Grant Number 203141/Z/16/Z. The OPIC electron microscopy facility was foundedby a Wellcome Trust JIF award (060208/Z/00/Z) and is supported by a Wellcome Trust equipment grant (093305/Z/10/Z). We are grateful for technical assistance from the OPIC staff.

