## Supplemental Data for "Production and characterisation of stabilised PV-3 virus-like particles using *Pichia pastoris*"

### Supplementary Information

**Supplementary Table 1.** Cryo-EM data collection and image processing for PV-3 SC8

| **Parameter** | **Value** |
|---|---|
| **Data Collection** | |
| Voltage (kV) | 300 |
| Magnification (×) | 60975 |
| Defocus range (µm) | -2.4 to -0.9 |
| Dose rate ($e^{-}$/pixel/s) | 9.20 |
| Frames | 20 |
| Frame length (s) | 0.150 |
| Total electron dose ($e^{-}$/Å$^2$) | 35.06 |
| Micrographs | 3717 |
| **Data processing** | |
| Pixel size (Å) | 0.82 |
| Initial particles (no.) | 10744 |
| Final particles (no.) | 2712 |
| Box size (pixels) | 450 |
| Symmetry | I1 |
| Accuracy of rotations (°) | 0.1320 |
| Accuracy of translations (Å) | 0.2662 |
| Resolution (Å) | 2.7 |
| Map sharpening *B*-factor (Å$^2$) | -59.1 |

**Supplementary Table 2.** Structure refinement and validation for the PV-3 SC8 capsid protein

| Parameter | Value |
|---|---|
| **Model composition** | |
| Non-hydrogen atoms | 5652 |
| Protein residues | 712 |
| Ligands | Sphingosine:1 |
| **Refinement** | |
| Resolution (Å) | 2.7 |
| Map CC[a] (Mask) | 0.85 |
| Map CC[a] (Volume) | 0.81 |
| **RMS deviations** | |
| Bond lengths (Å) | 0.002 |
| Bond angles (°) | 0.502 |
| **Mean B-factor (Å$^2$)** | |
| Protein | 13.05 |
| Ligand | 7.92 |
| **Validation** | |
| Molprobity[b] score (percentile) | 1.22 (99th) |
| Clashscore[b], all atoms (percentile) | 3.67 (97th) |
| Ramachandran favoured (%) | 97.71 |
| Ramachandran allowed (%) | 2.29 |
| Ramachandran outliers (%) | 0.00 |
| Rotamer favoured (outliers) (%) | 95.03 (0.32) |
| Cβ deviations >0.25 Å (%) | 0.00 |
| CaBLAM outliers (%) | 1.20 |
| CA Geometry outliers (%) | 0.44 |
| **EMRinger[c] score** | 4.63 |

[a] Map CC is given for the full particle reconstruction.

[b] Chen *et al.* (2010) Acta Crystallographica D66:12-21.

[c] Barad *et al.* (2015) Nature Methods 12:943–946.

**Supplementary Figure 1. Resolution of the PV-3 SC8 cryo-EM reconstruction. A** Fourier shell correlation (FSC) calculated between two independent half sets of data as a function of spatial frequency is plotted for the PV-3 SC8 cryo-EM reconstruction. FSC is plotted for the original unmasked half-maps (grey) and masked half-maps that had density corresponding to solvent removed (blue). FSC is also shown for phase-randomized half-maps (red) used to compensate for possible effects of the masking procedure before calculating the final corrected FSC (black). Good agreement between the masked and corrected curves indicated no adverse effects from the masking. The resolution at which the corrected curve drops below the FSC=0.143 threshold (grey dashed line) is indicated with an arrow. **B** Local resolution analysis of the final cryo-EM electron potential map for PV-3 SC8 as assessed by RELION local resolution estimation. A central slice through the VLP is viewed along the 2-fold axis and the distribution of local resolution is shown coloured from blue to red according to the colour key shown.

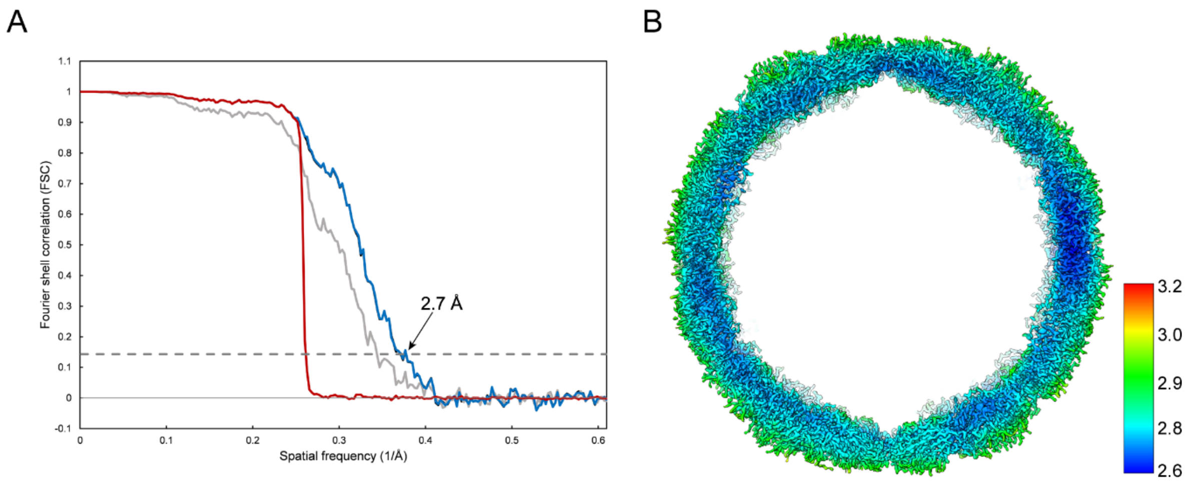
